# Inferring the genetic basis of sex determination from the genome of a dioecious nightshade

**DOI:** 10.1101/2020.07.23.218370

**Authors:** Meng Wu, Gregory J. Anderson, Matthew W. Hahn, Leonie C. Moyle, Rafael F. Guerrero

## Abstract

Dissecting the genetic mechanisms underlying dioecy (*i*.*e*. separate female and male individuals) is critical for understanding the evolution of this pervasive reproductive strategy. Nonetheless, the genetic basis of sex determination remains unclear in many cases, especially in systems where dioecy has arisen recently. Within the economically important plant genus *Solanum* (∼2000 species), dioecy is thought to have evolved independently at least 4 times across roughly 20 species. Here, we generate the first genome sequence of a dioecious *Solanum* and use it to ascertain the genetic basis of sex determination in this species. We *de novo* assembled and annotated the genome of *S. appendiculatum* (assembly size: ∼750 Mb; scaffold N50: 0.92 Mb; ∼35,000 genes), identified sex-specific sequences and their locations in the genome, and inferred that males in this species are the heterogametic sex. We also analyzed gene expression patterns in floral tissues of males and females, finding ∼100 genes that are differentially expressed between the sexes. These analyses, together with observed patterns of gene-family evolution specific to *S. appendiculatum*, consistently implicate a suite of genes from the regulatory network controlling pectin degradation and modification in the expression of sex. Furthermore, the genome of a species with a relatively young sex determination system provides the foundational resources for future studies on the independent evolution of dioecy in this speciose clade.

## INTRODUCTION

Dioecy—the presence of separate female and male individuals in a species, usually accompanied by dramatic phenotypic differences between the sexes—is a substantial, evolutionarily potent source of intraspecific variation. In species with genetic sex determination, these differences between males and females extend to their genomes, which show genomic differentiation in and around the sex determining region (or SDR) (Charlesworth 2002; Charlesworth 2012). Theory for the origin and emergence of these genomic signatures of dioecy—including sequence divergence in the SDR (Charlesworth and Charlesworth 1978; Rice 1987)—is well developed. According to these models, genetically determined dioecy starts with mutations at two loci, one mutation resulting in male sterility and one mutation leading to female sterility; these two mutations are expected to be tightly linked within a single genomic region. The SDR is later formed by the suppression of recombination between these two loci. After establishment, the SDR can continue to expand (i.e., encroach on neighboring genomic regions), increasing the amount of the genome that is differentiated between the sexes (Otto, et al. 2011). While some evidence is consistent with this “two-locus” model for the evolution of dioecy (see below), this pathway may not be the only way to achieve genetic sex determination (Henry, et al. 2018). Accordingly, it remains unclear whether the origin and early evolution of dioecy unfolds in predictable and generalizable ways across independent instances of the transition from hermaphroditism to dioecy.

Empirical data from a diversity of dioecious systems can provide critical information for understanding the underpinnings of this transition, and for identifying common genomic features of genetic sex determination. However, while dioecy is extremely common in animal species (where it is more often termed “gonochorism”), its origins are due to a few ancient events that are shared across major taxa, making analyses of the early emergence and evolution of dioecy challenging in animal systems. In contrast, while only 5-6% of angiosperm species have separate male and female individuals (Renner 2014), dioecy is phylogenetically widespread, occurring across 43% of angiosperm families (Renner 2014). It has been estimated that dioecy arose at least 100 times independently within the flowering plants (Charlesworth 2002; Renner 2014). This repeated emergence of dioecy in plants makes them ideal for the study of genetic sex determination and the early stages of sex-chromosome evolution, in ways that ancient and static animal systems cannot match (Charlesworth 2015).

Indeed, recent genomic analyses in young dioecious plant systems have made substantial progress toward evaluating the nature of the emerging SDR and uncovering the genetic factors and molecular mechanisms that regulate sex-determination. For instance, sex differences in frequency of *k*-mers (sequence motifs of varying lengths) have been used to identify sex-specific genomic sequences and to identify the SDR in persimmon, kiwifruit, and date palm tree (Akagi, et al. 2014; Akagi, et al. 2018; Torres, et al. 2018). Genomic approaches, combined with data from more classical analyses, have been used several times to test the two-locus model for the evolution of dioecy in plants (Charlesworth and Charlesworth 1978), with evidence for this model found in species such as *Silene*, papaya, asparagus, and kiwifruit (Wang, et al. 2012; Kazama, et al. 2016; Harkess, et al. 2017; Akagi, et al. 2019). In other species, however, a single gene seems to be sufficient for the expression of maleness and the repression of female development (as in persimmon) (Akagi, et al. 2014; Akagi, et al. 2016).

Despite this emerging clarity, several challenges still arise in studying the genetic basis of dioecy in angiosperms. Because dioecy in many plants is recently evolved, locating the SDR based on obvious signatures (such as large-scale genomic differentiation) used for many animal sex chromosomes (Charlesworth 2012, 2015) is often not appropriate. Compounding this problem, plant genomes are often rich in transposable elements, which can contribute to the fast accumulation of repetitive sequence around the non-recombining SDR (Li, et al. 2016); this can both obscure simple signatures of genomic differentiation expected to be associated with this region and can lead to difficulties in the *de novo* assembly and reconstruction of this region. Other challenges are biological. For instance, the genetic mechanisms involved in flower development are numerous and complex (involving various genes, small RNAs, epigenetic effects, and environment interactions) (Martin, et al. 2009; Nag and Jack 2010; Adam, et al. 2011; Song, et al. 2013; Akagi, et al. 2016; Bräutigam, et al. 2017; Murase, et al. 2017), making it difficult to pinpoint the specific loci responsible for sex determination in any particular case. In addition, while several studies have made important contributions to understanding the genetic basis underpinning the evolution of dioecy, most currently examined plant systems also have high levels of sexual dimorphism, amplifying the potential for multiple correlated phenotypic changes to confound or mask the causal loci involved in the initial emergence of sex differentiation.

Dioecious species in the genus *Solanum* present the opportunity to study the recent and repeated evolution of dioecy in this speciose and economically important plant clade. While most of the nearly 2000 species in the genus are hermaphroditic, there are a number of cases of dioecy, andromonoecy (in which male and hermaphrodite flowers occur separately on the same plant), and even sexually fluid species (Anderson 1979; Symon 1979; Symon 1981; Anderson and Levine 1982; Levine and Anderson 1986; Whalen and Costich 1986; Anderson and Symon 1989; Anderson, et al. 2015; McDonnell, et al. 2019). Of these, there are 15-20 clearly dioecious *Solanum* species (Anderson, et al. 2015), and their distribution across the phylogeny of the genus suggests that dioecy has independently evolved at least four times (that is, dioecious species appear in four different sections: one in the potato group, one in Dulcamaroid, two in Geminata, and 14 in Leptostemonum (Knapp, et al. 2004; Anderson, et al. 2015)). Interestingly, in all the dioecious species, female flowers still produce anthers, and the majority of the anthers on the female flowers also include pollen (Anderson, et al. 2015). In spite of high viability (stainability) of that pollen, and the presence of many of the same internal constituents as the pollen from male flowers, the pollen grains do not function in the usual way. They lack the typical three apertures (important for emergence of the pollen tubes from the pollen grains); i.e., they are inaperturate (Levine and Anderson 1986) (Figure 1). Nonetheless, the pollen in these female flowers plays a key role, i.e., is ‘functional’ in the other key role for pollen in *Solanum* flowers: there is no floral nectar (Anderson and Symon 1985), thus the inaperturate pollen is the only reward for the bee pollinators. Moreover, this consistent transition to inaperaturate pollen in female flowers suggests that independent transitions to dioecy within *Solanum* might often involve convergent phenotypic changes, at least for the emergence of females. Furthermore, there are currently no genomic analyses of dioecious *Solanum* species, and nothing is known of the mechanistic genetic basis of dioecy in this genus.

**Figure 1.**
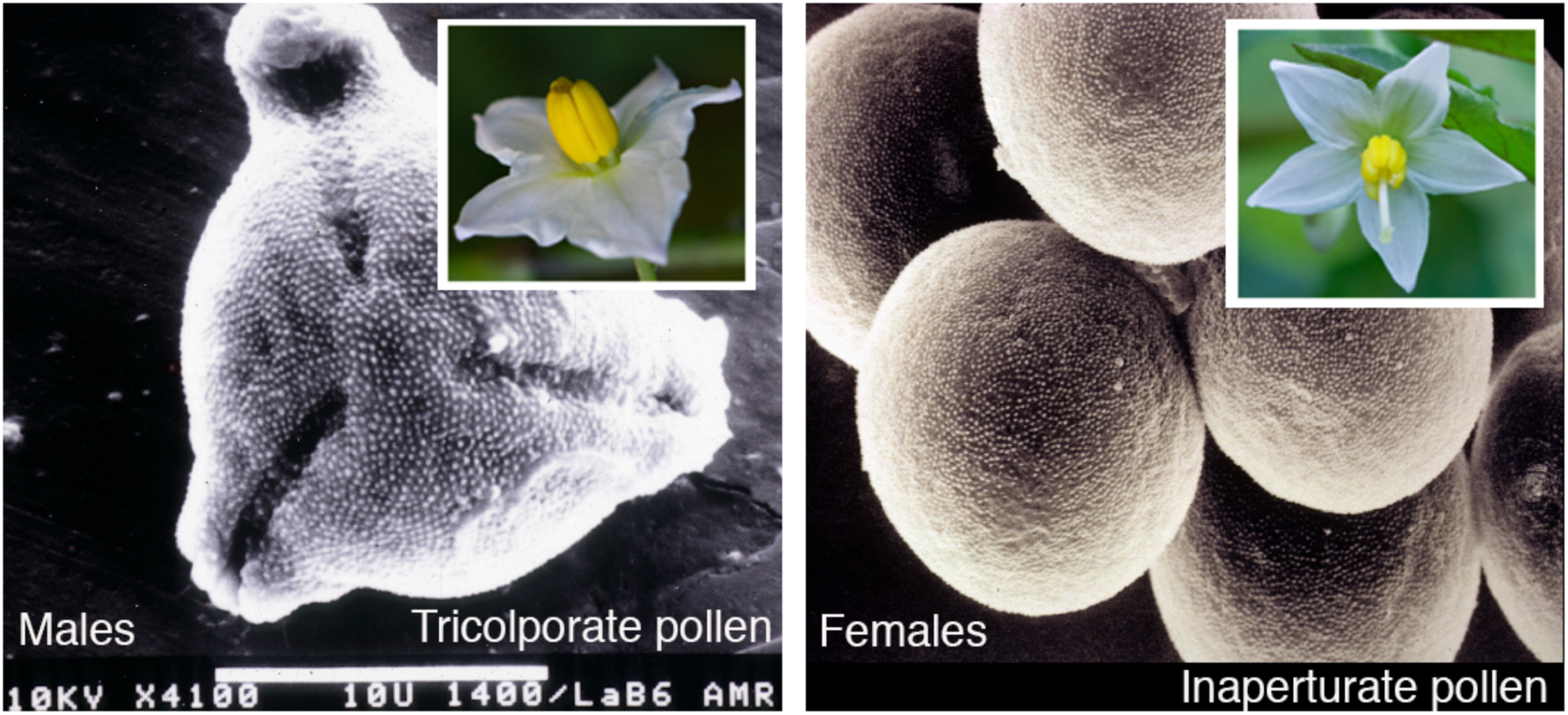
Pollen produced in male and female flowers of *S. appendiculatum*. On the left, male pollen is tricolporate, typical of the genus. On the right, inaperturate pollen produced by females is shown (lacking pores). Flowers of each sex are shown in the upper inset.

Thus, our goal was to address the mechanisms underpinning the emergence of dioecy in one such *Solanum* species. *Solanum appendiculatum* is the sole dioecious species in the Anarrhichomenum section of *Solanum*, which is estimated to be only 4 million years old (Echeverría-Londoño, et al. 2020). There is no evidence of sex chromosome divergence in the karyotypes of *S. appendiculatum* (Bernardello and Anderson 1990), and it is unknown whether males or females are heterokaryotypic (i.e., whether this is a XX/XY or ZZ/ZW sex-determination system). Consistent with a very recent origin of dioecy (<4 MY), this species shows very little sexual dimorphism: female flowers are morphologically hermaphroditic but functionally pistillate due to the non-germinable inaperturate pollen, while the male flowers never set fruit (Figure 1) (Anderson 1979). Previous studies of the early development of pollen in *S. appendiculatum* showed that pollen from the female flowers develops a primexine (a cellulose layer that initiates the pollen wall formation) at the physical position that would otherwise develop apertures (Zavada and Anderson 1997). Intine (consisting primarily of cellulose and pectin) then accumulates and thickens the pollen wall at these locations, resulting in the failure to develop germination pores (Zavada and Anderson 1997). Although the sexual system of this species is well understood, including the structural basis of functional male sterility in females, the genetic basis of sex determination is unknown.

In this study, we generated a high-coverage genome assembly of *S. appendiculatum* by adopting a hybrid assembly strategy using PacBio long reads and Illumina short reads. We paired this with RNA-seq data from male and female bud and flower tissue. Based on the genome annotation in this new assembly, we assessed three lines of data for genomic signatures of sex-determination: a) we investigated gene family dynamics through comparative analyses with seven other annotated (non-dioecious) genomes in the Solanaceae; b) we identified sex-biased gene expression in floral bud tissue and mature flowers; and, c) we identified candidate sex-specific regions based on the location of sex-specific *k*-mers in the genome assembly. Interestingly, we uncovered pectin-related genes in all of these analyses. Pectin is associated with several sex-specific functions in plants (Micheli 2001), including a direct functional role for pectin production in pollen wall and germination pore development (Gou, et al. 2012; Jiang, et al. 2013). Therefore, we propose that the putative mechanism of the transition to inaperturate pollen in females of *S. appendiculatum* is a change in pectin regulation in female flowers. This genome assembly, and the putative SDR and candidate loci underpinning sex determination that emerge from it, also provide a clear framework for future studies of sex determination more broadly in *Solanum*.

## MATERIALS AND METHODS

### Plant sampling and sequencing

Individuals from two accessions (population locations) of *S. appendiculatum* (denoted accessions #670 and #716 by G. Anderson), originally collected from natural habitats in Mexico, were grown and maintained in the greenhouse at Indiana University. The original collection data and voucher information for these accessions are provided in the supplementary materials. We collected young leaves from male and female individuals (three of each from accession 670, two of each from accession 716) and extracted DNA using the Qiagen DNeasy Plant Mini Kit. We treated samples with RNAase-A and used 100 uL of elution buffer. From each of these samples, we sequenced genomic DNA using the Illumina sequencing-by-synthesis technology (2×150 bp using HiSeq). Additionally, we obtained long-read sequences from one male and one female (670-1190 and 670-34) using the PacBio platform (sequencing carried out by Novogene Hong Kong).

We collected buds (1 day prior to anthesis) and mature flowers from 12 individuals (3 males and 3 females from each accession) on the same day. Additionally, we collected young leaves from the reference female (670-1190) from which we extracted RNA for genome annotation purposes. Tissues were ground using the Tissue-lyser (Qiagen), RNA was extracted using the Qiagen RNeasy Plant mini-kit and brought to a final concentration of 70–200 ng/uL. A subset of RNA samples was checked for quality by agarose gel electrophoresis and visualized by ethidium bromide staining. Sample quality was further evaluated using the Agilent 2200 RNA TapeStation system before library creation. Stranded, paired-end libraries of total RNA were generated for each sample using Illumina Truseq Stranded mRNA sample preparation kits. Short-read DNA and RNA quality control, library preparation, and pooling were performed by the Indiana University Center for Genomics and Bioinformatics.

### Genome assembly and quality assessment

We followed our previous workflow (Wu, et al. 2019) to assemble and annotate the genome. In brief, we adopted the MaSuRCA v3.2.2 (Zimin, et al. 2017) pipeline to generate the genome assembly, in which relatively high-accuracy Illumina paired-end reads from one male plant (121X coverage from 670-1190) and one female plant (116X coverage from 670-34) were used together to trim and correct low base-call-accuracy long-reads generated by PacBio sequencing from the same two plants (19X coverage from the male 670-1190 and 21X coverage from the female 670-1190). Genome size was estimated based on the *k*-mer abundance distribution separately from the male and female reads using GenomeScope (Vurture, et al. 2017), with a *k*-mer length of 25-bp and max *k*-mer coverage of 10,000. After initial assembly, all assembled scaffold sequences were aligned against bacterial, archaea, fungal, and human databases to remove potential contaminants in our assembly, using the DeconSeq tool v0.4.3 (Schmieder and Edwards 2011).

We evaluated the completeness of the genome assembly using 1,515 plant near-universal single-copy orthologs within BUSCO v3 (Simão, et al. 2015). To provide an estimate of assembly accuracy at the nucleotide level, we calculated a quality score for every position in the genome assembly using the program Referee (Thomas and Hahn 2019). Referee compares the log-ratio of the sum of genotype likelihoods for the genotypes that contain the reference base (e.g. [A, A], [A, T], [A, C], and [A, G] for reference base ‘A’) vs. the sum of those that do not contain the reference base (e.g. [T, T], [T, C], [T, G], [C, C], [C, G] and [G, G] for reference base ‘A’). The input used in the Referee calculation was obtained from the output pileup file from ANGSD (Korneliussen, et al. 2014), which pre-calculated genotype likelihoods at each base of the genome assembly. Here, two genotype likelihood scores for every position in the genome assembly were calculated separately based on either the BAM file of aligned Illumina reads from the male (770-34) or the BAM file of aligned the Illumina reads from the female (670-1190).

### Repeat and Gene Annotation

We followed the “Repeat Library Construction - Advanced” steps from the MAKER-P pipeline (Campbell, et al. 2014) to generate a species-specific repeat library. Briefly, the LTR retrotransposon (LTR-RT) library was constructed using LTR-harvest (Ellinghaus, et al. 2008) and LTR-retriever (Ou and Jiang 2018). Miniature inverted transposable elements (MITEs) were detected by MITE-Hunter (Han and Wessler 2010). Other repetitive sequences were identified with RepeatModeler (http://www.repeatmasker.org/RepeatModeler/). All the detected repetitive elements above were combined into the *S. appendiculatum*-specific library. RepeatMasker (Smit, et al. 2015) was then used to mask all repeat elements in the assembled genome by searching for homologous repeats in the species-specific library.

Gene models were predicted using three classes of evidence: RNA-seq data, protein homology, and *ab initio* gene prediction (see Supplementary text), which was performed within the MAKER-P pipeline (Campbell, et al. 2014). All information on gene structures from the three different classes was synthesized to produce final gene annotations. We extracted relatively high-confidence genes according to evidence-based quality, requiring an annotation evidence distance (AED, which measures the goodness of fit of an annotation to the RNA/protein-alignment evidence supporting it) score <0.5. To assign functions to genes, we followed the pipeline AHRD (https://github.com/groupschoof/AHRD) to automatically select the most concise, informative, and precise functional annotation (see Supplementary text). For each gene, the associated gene ontology (GO) annotations were assigned according to the predicted protein domains (http://www.geneontology.org/external2go/interpro2go). To investigate the putative genomic locations of predicted genes in the genome, we performed whole-genome synteny alignment using Satsuma v3.1.0 (Grabherr, et al. 2010), and recorded the genes unambiguously associated with a single identified syntenic region in the tomato (*S. lycopersicum*) genome (version ITAG3.20) (The Tomato Genome Consortium 2012).

### Gene family analyses

To investigate changes in gene family size specifically in *S. appendiculatum*, we determined the expanded or contracted gene families along this branch in a phylogenetic tree. We obtained the protein-coding sequences of several Solanaceae species with high-quality genome annotations, including *Solanum lycopersicum, Solanum tuberosum, Jaltomata sinuosa, Nicotiana attenuata, Petunia axillaris*, and *Petunia inflata*. Gene families from these species were inferred by the program OrthoFinder (Emms and Kelly 2015), as were the phylogenetic relationships among species. Divergence times were retrieved from previous estimates within the Solanaceae (Särkinen, et al. 2013). Rapid evolving gene families along the branch leading to *S. appendiculatum* were determined using the program CAFE v3.1 (Han, et al. 2013) with a *P*-value cutoff of 0.01.

### Gene expression analyses

For the RNA-seq data, trimmed reads from each library were mapped against the assembled genome using HISAT v2.1.0 (Kim, et al. 2015), and the resulting SAM files were then converted to sorted BAM files using SAMtools v0.1.19 (Li, et al. 2009). Reads assigned to exonic regions of genes were counted, and read counts of each annotated gene were calculated using FeatureCounts (Liao, et al. 2013). The raw read count table was used as input for the program edgeR (Robinson, et al. 2010). For downstream analysis, we removed genes with low expression by requiring each gene to have >1 read count per million (CPM) mapped reads in at least two samples (of any accession, sex, or tissue type). Trimmed mean of *m-*value (TMM) normalization was performed to eliminate composition biases between different libraries (Robinson and Oshlack 2010). Based on gene expression patterns, we also performed multi-dimensional scaling analysis in edgeR (Robinson, et al. 2010) using the function “PlotMDS.” The distances among samples were estimated using the average of the absolute value of the log2-fold expression changes among genes.

Differential expression analysis was performed separately for flower buds and mature flowers using edgeR (Robinson, et al. 2010). We fit an additive model (expression ∼ accession + sex) to control for the underlying differences among accessions, using the *glmfit* function. The genes that consistently displayed differential expression for a given pairwise sex comparison were identified using the function *glmLRT*. Genes were only considered to be significantly differentially expressed between the two sexes at a false discovery rate (FDR) < 0.05 (Benjamini and Hochberg 1995) and fold change (FC) >2.

### Search for putative sex-determination region

We used a *k*-mer method to find putative sex-determination region sequences (Akagi, et al. 2014; Akagi, et al. 2018). First, we identified putative sex-specific regions by examining different coverage of male/female reads across genome. We defined genomic regions of interest by identifying overlapping 10-Kb windows that had high counts of sex-specific reads (defined as more than 2 counts per million), in which more than 30% of the window was covered by the sex-specific reads, and for which sequence divergence among the sexes was larger than among populations.

Second, we defined *k*-mers as sex-specific if they were detected in all samples of one sex but were completely absent in the samples of the other. We first extracted all 30-bp *k*-mers from the genomic short reads of six males and females (the same dataset used in variant calling), and counted the frequency of each *k*-mer in each sample using Jellyfish v2.2.9 (Marçais and Kingsford 2011). We determined the genomic location of sex-specific *k*-mers through the sequence alignment information of read pairs that contained them (from the mapping used for variant calling). Counts of female-specific and male-specific reads were calculated for non-overlapping 10-Kb window genome-wide using BEDTools (Quinlan and Hall 2010), which also calculates the breadth of the covered region (i.e. the proportion of a 10-Kb window being covered by at least one sex-specific read)(Quinlan and Hall 2010). We then used the counts and distribution of sex-specific *k*-mers across the 10-Kb windows to narrow down the candidate sex-specific regions. To investigate sequence divergence between two sexes, we calculated measures of population differentiation (*d*_*XY*_ and *F*_ST_) along with genetic diversity (*π*) in 10-kb windows (using the python scripts at https://github.com/simonhmartin/genomics_general). We filtered out low-quality sites (genotype quality <30 or read depth <5 in each individual) and used only windows with at least 1000 sites after filtering. We calculated divergence among populations (*i*.*e*. accessions 670 and 716) and between males and females, to then extract the windows in which male-female *d*_*XY*_ was larger than that between the two populations.

## RESULTS

Following the workflow (Figure S1), we generated the first genome assembly of a dioecious species in *Solanum* (Table 1). The genome of *S. appendiculatum* is estimated to be 671.8 Mb based on the *k*-mer frequency distribution, which is consistent with the estimated size (750 ±14 Mb for females and 742 ±6 Mb for males; *n* = 3 for each sex) from flow cytometry (Haak DC, unpublished data). The size of the genome assembly is ∼751.9 Mb, which includes 3,643 scaffolds with an N50 length of ∼920.8 Kb. In the assembly, ∼96.6% of 1,515 BUSCO plant universal single-copy orthologous genes were found to be complete. 99.6% of sites in the assembly were well supported by either male or female Illumina reads with a quality score higher than 20 (Table S1), suggestive of low base-calling error in our assembly. Base calls in the assembly were slightly better supported by female reads than male reads (99.4% vs. 97.7%) at a quality threshold of 20 as determined by separate runs of the Referee program (Thomas and Hahn 2019) (Table S1).

**Table 1.** Summary of the *S. appendiculatum* Genome Assembly

|  |  |
| --- | --- |
| **Assembly features** |  |
| Genome size estimated (Mb) | 671.83 |
| Assembly size (Mb) | 751.93 |
| Number of scaffolds | 3643 |
| Scaffold N50 length (Kb) | 920.78 |
| GC contents | 35.78% |
| Plant_CEGs (BUSCO)<sup>a</sup> | 92.6+(4.0+1.0) |
| Bases with quality score >20 | 99.57% |
| **Repeat annotation** |  |
| Total (Mb) | 497.56 (66.17%) |
| Gypsy (Mb) | 254.87 (33.90%) |
| Copia (Mb) | 26.98 (3.59%) |
| Unknown LTR-RTs (Mb) | 90.79 (12.06%) |
| **Gene annotation** |  |
| Number of protein-coding genes | 35,731 |
| Genes supported by RNA-seq | 82.08% |
| Mean CDS length (bp) | 1265.56 |
| Number of exons per gene | 6.17 |

<sup>a</sup>Plant\_CEGs (Clusters of Essential Genes) shows the percentage of complete single-copy orthologs plus the percentage of duplicated orthologs and fragmented orthologs.

The genome of *S. appendiculatum* is highly repetitive, consisting of 497.6 Mb of repetitive sequence (66.2% of the assembly; Table S2). Consistent with other Solanaceae genomes, *Gypsy* LTR-RTs are the most abundant repeat elements in the *S. appendiculatum* genome (254.9 Mb; comprising 33.9% of the assembly). To annotate protein-coding genes we used the MAKER-P pipeline, finding 35,731 high-confidence genes with AED scores <0.5, which is similar to the gene number (35,768) annotated in the domesticated tomato genome (The Tomato Genome Consortium 2012). The gene structure of annotated genes (e.g., CDS lengths, exon numbers, and GC contents) were also similar to the genes annotated in other Solanaceae genomes (Table 1). Among high-confidence genes, we found 82.1% of their exonic regions are supported by at least one RNA-seq read, and 64.3% of them have expression levels with TPM>1 (i.e. transcript per million) in at least one sample of our investigated flower or leaf tissues. Together, the large scaffold size, the high coverage of the plant conserved single-copy genes, the low base-calling error, and the well-supported gene annotation all indicate a high-quality assembly for this species.

### Sex-biased gene expression is greatest in mature flowers

We tested for differential gene expression among sexes, separately in flower buds and in mature flowers. In buds, only 16 genes were identified to be significantly sex-biased (Figure 2A; Table S3), while 95 significantly sex-biased genes were detected in mature flowers (Figure 2B; Table S4). Almost two-thirds (58) of the differentially expressed genes in mature flowers are female-biased, with higher expression in the female flowers (Figure 2B; Table S4).

**Figure 2.**
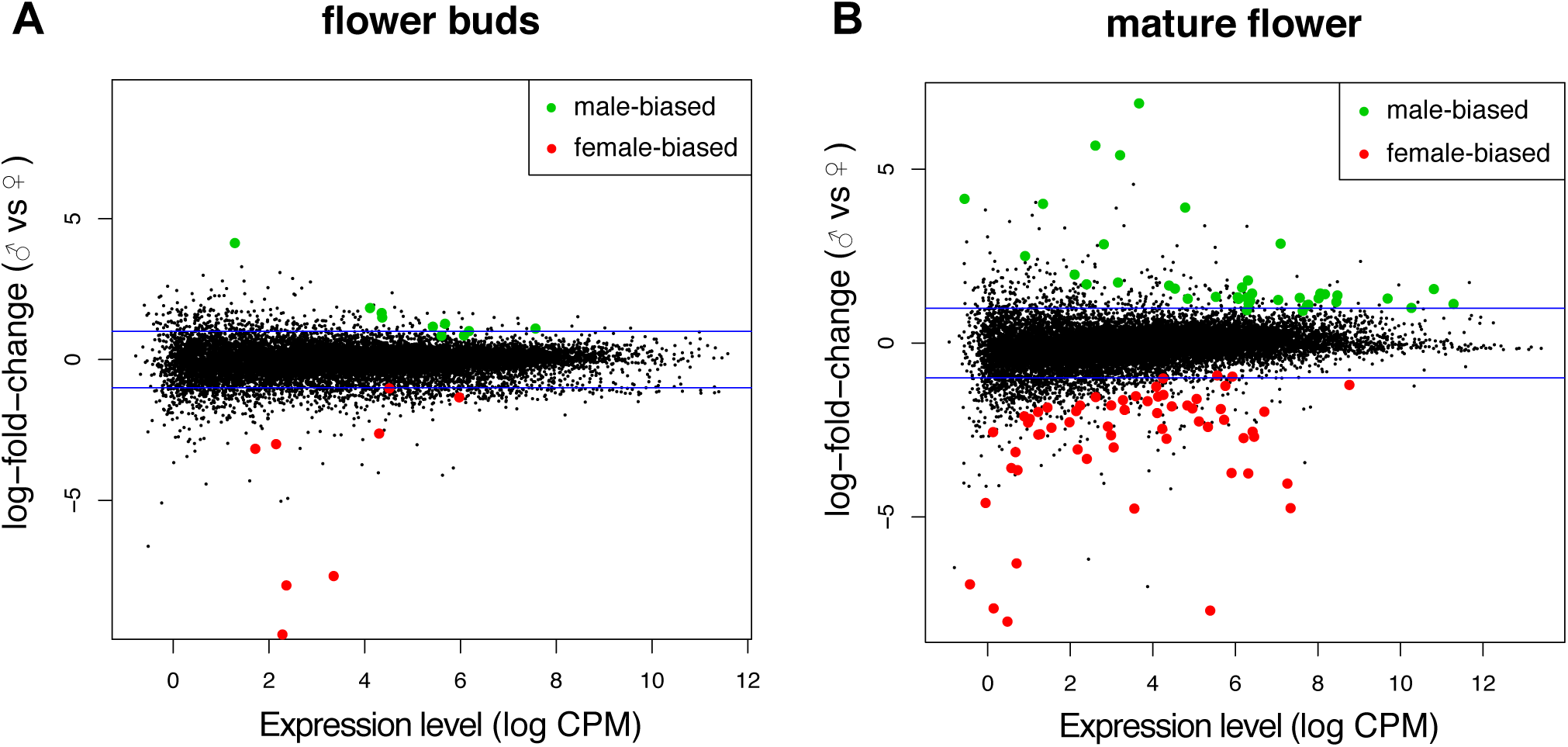
Sex-biased gene expression pattern at two different flower development stages of *S. appendiculatum*. Differentiation gene expression between male and female flowers in **(A)** flower buds, and **(B)** mature flowers.

Among the genes showing sex-biased expression in mature flowers, we found several that may be functionally associated with phenotypic differences between males and females. In particular, we identified eight female-biased genes that are functionally related to pectin metabolism, which is involved in the production of germination pores on pollen (see Discussion). These eight genes include genes encoding a pectin acetylesterase family protein, a pectin lyase-like protein, and an invertase/pectin methylesterase inhibitor. The frequency of pectin-related genes is significantly higher in the set of sex-biased genes (Fisher’s Exact *P* = 3.17^-5^) and female-biased genes (Fisher’s Exact *P* = 1.05^-6^), compared to pectin-related genes as a fraction of all genes (416 out of 35,731 annotated genes). Several of the female-biased genes also appear to be pistil-specific (that is, found only in female reproductive tract or ovary: the ‘pistil’). In particular, the gene *SlINO* has female-biased expression, and is a known pistil-specific *YABBY* transcription factor in domesticated tomato (Ezura, et al. 2017).

### Candidate sex-determination regions suggest an XX/XY system

Based on a search for sex-specific sequences in six male and six female individuals, we identified ∼11,000 female-specific and ∼6,000 male-specific *k*-mers. The relatively few sex-specific sequences (compared to other species) may indicate a very recent origin of dioecy, a very small sex-determining region, or both. We found 18 windows enriched with sex-specific reads, out of which seven (on four scaffolds: scf15976, scf14997, scf15476 and scf16572) have significantly elevated sequence divergence (*d*_*XY*_) between males and females (Figure 3). Six out of these seven candidate windows are enriched for male-specific sequences, and therefore likely represent male-specific regions. Heterozygosity was found to be higher in males in all of these male-specific regions, as expected in an XX/XY sex-determination system. Moreover, two of these male scaffolds (scf14997 and scf16572) are orthologous to regions just 1 Mb apart on chromosome 12 of *S. lycopersicum* (genome version ITAG3.20; Figure S2), indicating that they may be one region in *S. appendiculatum*. Together, these results suggest that *S. appendiculatum* has an XX/XY sex determination system. While we found more female-specific *k*-mers relative to male-specific ones, the differences in sequencing coverage among sexes (Table S8) suggest that this observation should not be interpreted as evidence supporting a ZZ/ZW system.

**Figure 3.**
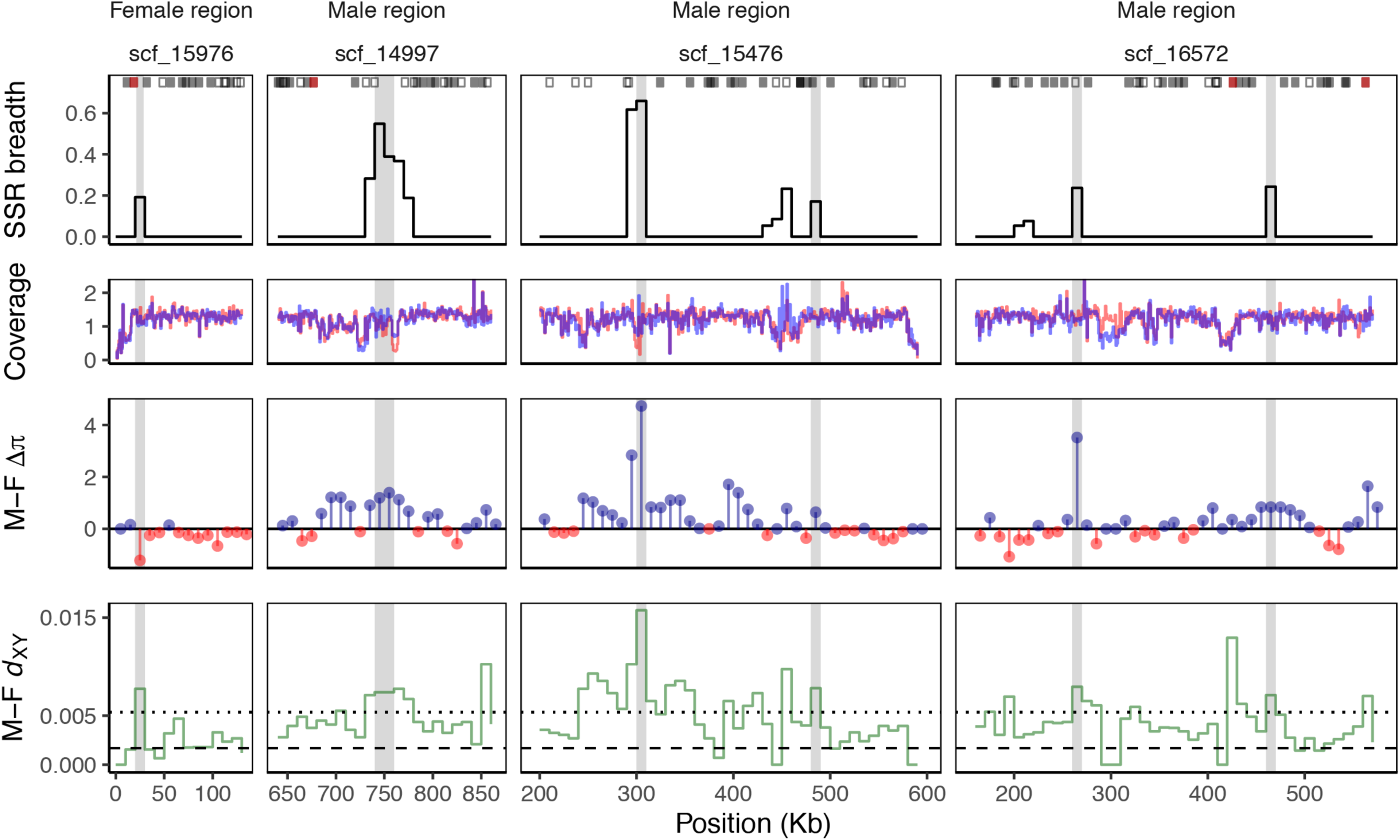
Four genomic scaffolds that contain seven 10-Kb windows enriched with sex-specific sequences (shaded light grey areas). Top row: Sex-specific read breadth (fraction of window covered by reads with sex-specific sequence). Squares are annotated genes in each scaffold (red: differentially expressed in at least one floral tissue; black: expressed in floral tissue but with no differential expression between sexes; open: not expressed in floral tissue). Second row: Normalized read depth in male (blue) and female samples (red). Third row: Difference in heterozygosity level (π) between males and females. Positive values indicate higher sequence diversity in males. Last row: Absolute sequence divergence (*d*_XY_) between males and females.

We annotated 20 genes within male-specific regions and 5 genes in the sole female-specific scaffold (Table S9). Several of these annotated genes have been previously implicated in flower development and sex determination. Notably, we found three invertase/pectin methylesterase inhibitors (PMEIs; genes *sapp25116, sapp25117* and *sapp25118* on scaf14997), which could play a role in regulating pectin degradation or modification in pollen grains. The three PMEIs have sequence identity >90% and are located next to each other within a single syntenic block, consistent with recent tandem duplication events. Other genes found in these sex-specific regions are potentially involved in pollen grain cell division, ethylene signaling, and anther development (all summarized in Table S9).

### Pectin-related gene families have evolved dynamically in S. appendiculatum

We identified 228 rapidly expanded gene families and 75 rapidly contracted gene families on the branch leading to *S. appendiculatum* (Table S5; Table S6). As expected from previous studies in plants, many of those rapidly evolving gene families are involved in stress-related response, such as NBS-LRR disease resistance proteins. Of the 228 rapidly expanding gene families in *S. appendiculatum*, we found five gene families that are functionally related to pectin degradation and modification. These included genes in the rhamnogalacturonate lyase, pectin lyase, invertase/pectin methylesterase inhibitor, and pectin-acetylesterase families (Table S7). In this last expanded family, there are two female-biased genes (*sapp52355* and *sapp52361*), as identified in our RNA-seq data (above). Lineage-specific expansion of pectin-related gene families was also found in other investigated Solanaceae genomes, but a smaller number of genes were duplicated in those species relative to *S. appendiculatum* (Table S7). In addition to pectin-related genes, we also detected two rapidly expanded gene families (OG0001468 [+3] and OG0001481 [+5]) and one rapidly contracted gene family (OG0000179 [-3]) annotated as “plant self-incompatibility protein S1 family”.

## DISCUSSION

We generated the first genome for a dioecious species within the genus *Solanum*, to assess the early emergence and genomic signatures of sex-differentiation and sex-determination. To do so, we assembled a high-quality genome, took a *k*-mer approach to find sex-linked genomic regions, and carried out an RNA-seq experiment of floral tissues to find genes involved in sex determination and sexual dimorphism. We found that dioecious *S. appendiculatum* appears to have a very small, recently evolved sex-determination system and that males are likely to be the heterogametic sex. Moreover, the specific loci associated with sex-differentiation suggest that the evolution of dioecy in this system involved changes in the regulation of pectin synthesis and degradation, including in specific phenotypic transitions observed in functionally female flowers. This genome, and the associated candidate genes, represents a valuable genomic resource for the continued investigation of recent transitions to dioecy within *Solanum*.

### Limited sex-biased gene expression and few sex-associated regions are consistent with recent evolution of sexual dimorphism

We found a very modest amount of sex-biased gene expression in flower buds, and larger but still delimited sex differences in the expression profiles of mature flowers. Given that sex-specificity of gene expression is expected to accumulate with time since the origin of sexual dimorphism (Ellegren and Parsch 2007), the observation that few genes show sex-biased expression is consistent with a young sex-determination system. This very modest genomic and transcriptomic divergence between the sexes is consistent with the subtle morphological differentiation between male and female flowers, which is among the least pronounced in the dioecious nightshades (Anderson, et al. 2015).

For mature flowers, sex-biased genes more commonly had higher expression in females than in males (Figure 2B). This finding contrasts with another species with a recently evolved sex-determining region--the garden asparagus (Harkess, et al. 2015)—likely because of developmental differences in sex expression between the two systems. In asparagus, anther development is arrested before microspore meiosis in female flowers (Caporali, et al. 1994), thus genes associated with later pollen development are expected to be expressed only in males (Harkess, et al. 2015). In contrast, in *S. appendiculatum* female flowers develop mature pollen, but fail to deposit primexine at the apertural regions (Zavada and Anderson 1997). Our observation of more female-biased genes in *S. appendiculatum* is therefore consistent with this maintenance of both functional styles (female reproductive parts) and active production of (inaperturate) pollen (Levine and Anderson 1986) in female flowers, and seems to indicate some loss of function of female reproductive parts in male plants. This possible loss of function, however, is not reflected in the morphology of male flowers, which have complete female reproductive parts (albeit with much shorter styles; (Anderson 1979; Anderson and Levine 1982)).

### Regulation of pectin as a potential mechanism for the formation of aperturate pollen

Identification of candidate genes playing potential feminizing or masculinizing effects is important to understand sex determination in this recently evolved dioecious species. Collectively, three different approaches in this study—gene family dynamics, sex-biased expression, and sex-specific *k*-mers –detected a set of loci distinctive to *S. appendiculatum*. Some of these are likely unrelated to this species’ transition to dioecy, and some others are possibly associated with general physiological consequences of this breeding system transition rather than directly involved in sex-differentiation and sex-determination *per se*. For instance, our gene family analysis detected a contraction of the self-incompatibility protein S1 family specifically in *S. appendiculatum*. Because the evolution of dioecy obviates the possibility of self-fertilization, this transition might be expected to relax selection to maintain functional self-incompatibility genes; similar losses of self-incompatibility proteins has also been observed in other Solanaceae species that have undergone breeding system transitions (e.g. to self-compatibility (Wu et al. 2019)). Nonetheless, among the genetic changes detected, it is striking that all three of our different approaches detected pectin-related genes in association with sexual differentiation in *S. appendiculatum*, including pectin acetylesterases (PAE), pectin lyase-like proteins (PLL), and pectin methylesterase inhibitors (PMEI). Our finding is particularly intriguing as pectin synthesis and regulation is known to play important roles in pollen wall development, and in pollen function more broadly. Pectin consists of homogalacturonan (HG), which can be methyl- and acetyl-esterified (Wu, et al. 2018), and pectin polysaccharides are critical components of the pollen wall. Mutants in genes encoding pectin polysaccharide synthetic and degrative enzymes—including pectin methylesterase (PME), polygalatcturonase (PG), PAE, and PLL—often show defective primexine, intine, or other pollen wall structures (Shi, et al. 2015; Wu, et al. 2018). Strikingly, in *Nicotiana* (Solanaceae), transgenic mutants of one pectin acetylesterase gene, *PAE1*, exhibit the loss of germination pores on the surface of the pollen grains (Gou, et al. 2012)—a very similar phenotype to the inaperturate pollen observed in the female flowers of *S. appendiculatum*. The overexpression *PAE1* in transgenic tobacco results in severe male sterility by affecting the germination of pollen grains and the growth of pollen tubes (Gou, et al. 2012).

Other pectin associated proteins are also implicated in numerous functional roles in pollen tube germination and growth, including via coordinated regulation between PMEs and their inhibitors—PMEIs (Mollet, et al. 2013). For instance, PME is important for the generation of methyl esterified HG in the apical zone of growing pollen tubes, which provides sufficient plasticity for sustaining growth (Cheung and Wu 2008). The removal of methyl ester groups by PME may allow the pectin-degrading enzymes, such as PLL or PG, to cleave the HG backbone, which may affect the rigidity of the cell wall (Gaffe, et al. 1994; Micheli 2001). It has been proposed that the pollen cell might maintain a closely regulated level of PME activity, via regulation by PMEIs, in order to maintain the equilibrium between strength and plasticity in the apical cell wall (Bosch and Hepler 2005, 2006). For example, silencing of the *PME1* gene in tobacco (Bosch and Hepler 2006), and suppression of PMEI *At1g10770* in *Arabidopsis* (Zhang, et al. 2010), both result in slowed pollen tube growth.

Interestingly, in addition to detecting sex-specific expression of PAE, we also found three PMEIs in a candidate sex-determining region (scf14997) in *S. appendiculatum*. The arrangement and relationship between these putative sex-determining genes is consistent with them being recent duplications, similar to what has been found in other dioecious plants (Harkess, et al. 2017; Akagi, et al. 2018). While the specific function of these genes is not yet known, the general roles of PMEIs, PAE, and other related proteins in the formation and function of pollen suggests some possible models for the emergence of sex-specific pollen functions in the two sexes of *S. appendiculatum*. For example, it is possible that these PMEI copies influence the differential (sex-specific) expression patterns of downstream pectin-related genes in mature flowers, including PAE, thereby inhibiting or initiating the feminizing effect (i.e. inaperturate pollen) observed in female flowers. This process could also involve other tightly linked genes: the same syntenic block contains a gene coding for a *LOB* domain protein (*sapp25115*), the *Arabidopsis* ortholog of which (*AT1G06280*) is specifically expressed during tapetum and microspore development in the anthers (Oh, et al. 2010; Zhu, et al. 2010). Other differentially expressed genes also have clearly relevant functions. For example, the pyruvate dehydrogenase E1 component subunit alpha (*sapp29734*) was differentially expressed between males and females in the mature flower; pyruvate dehydrogenase catalyzes the early steps of sporopollenin biosynthesis, a major component of the exine layer of pollen grains (Jiang, et al. 2013).

While pectin-related genes are promising candidates for the expected male-sterilizing step in the evolution of dioecy, it is possible that they are downstream of a master regulator of sex determination. Whether some upstream genetic changes trigger the downstream changes in pectin-related genes can be addressed in future studies. For instance, transcriptome analysis of additional developmental stages of male and female flowers could clarify how pectin regulation changes across flower development and the specific timing of divergent expression differences between male and female flowers. Regardless, with a genome-wide search for sex-specific sequences, in conjunction with gene expression analyses, we were able to detect both putative sex-determining regions and genes that may contribute to at least one of the two steps expected in the path from hermaphroditism to dioecy. These loci provide clear candidates for direct functional analysis in this system, especially for inaperturate pollen development phenotypes in female flowers.

### The S. appendiculatum genome provides a foundation for addressing repeated transitions to dioecy

Although the economically important, highly speciose (∼2000 species) plant genus *Solanum* contains fewer than 20 documented dioecious species, dioecy is estimated to have arisen independently at least 4 times (Anderson, et al. 2015). Many of these transitions appear to involve common phenotypic features, most notably the development of inaperturate pollen in female individuals and dramatic reduction of the pistil in male flowers (Anderson, et al. 2015). As such, this young genus (estimated ∼17MY old;(Särkinen, et al. 2013)) offers a promising system in which to address the genomic features and genetic mechanisms of repeated, recent transitions to dioecy.

*Solanum appendiculatum* is among the most recently evolved dioecious angiosperms with sequenced genomes (<4MY; (Echeverría-Londoño, et al. 2020)). As such, the resources generated here provide a valuable framework for examining additional transitions to dioecy in the highly speciose genus, including a high-quality assembled genome, transcriptome characterization for annotation and gene expression analyses, and a set of candidate loci for directed exploration in parallel systems. Because most dioecious nightshades have similar sexual traits, including inaperturate pollen in the stamens of female flowers (Anderson, et al. 2015), addressing the parallel origins of dioecy in this group can also address whether these transitions have followed convergent paths at genomic, genetic, and developmental levels. In conjunction with the *S. appendiculatum* genome, sequence data from other dioecious *Solanum* species can be used to dissect these parallel origins of sex determination in *Solanum*, including whether these exhibit similar genomic features (in terms of the number, size, and distribution of emerging sex-determination regions), draw on the same kinds of genomic/genetic changes (i.e. share orthologous sex-linked regions), and/or involve the same specific pathways and individual loci, including whether there is a general role for pectin-related loci in the early emergence of sexual differentiation. In this context, study of the genetic control of sex expression in species like S. *polygamum* and *S. conocarpum*—both of which bear anthers on female flowers, but that anthers are largely devoid of any pollen (Anderson et al. 2015)—could prove especially informative. Data from multiple recent, parallel systems will also be critical for testing the general predictions of theoretical models of the evolution of dioecy and assessing whether the complexity of genomic transitions that underpinning real empirical transitions matches well with these theoretical expectations.

## ACKNOWLEDGEMENTS

We are grateful to JL Kostyun and DC Haak for their help in the lab and greenhouse. This work was done with support from NSF grant IOS-1127059 and the Department of Biology at Indiana University. Previous support from NSF, and the University of Connecticut, supported the field collections and greenhouse experiments with *S. appendiculatum*.

## DATA DEPOSITION

The project has been deposited at NCBI BioProject database under the accession number PRJNA561636.

## Notes

### Competing Interest Statement

The authors have declared no competing interest.

## References

Adam H, Collin M, Richaud F, Beulé T, Cros D, Omoré A, Nodichao L, Nouy B, Tregear JW. 2011. Environmental regulation of sex determination in oil palm: current knowledge and insights from other species. Annals of botany 108:1529–1537.

Akagi T, Henry IM, Kawai T, Comai L, Tao R. 2016. Epigenetic regulation of the sex determination gene MeGI in polyploid persimmon. The Plant Cell 28:2905–2915.

Akagi T, Henry IM, Ohtani H, Morimoto T, Beppu K, Kataoka I, Tao R. 2018. A Y-encoded suppressor of feminization arose via lineage-specific duplication of a cytokinin response regulator in kiwifruit. The Plant Cell:tpc. 00787.02017.

Akagi T, Henry IM, Tao R, Comai L. 2014. A Y-chromosome–encoded small RNA acts as a sex determinant in persimmons. Science 346:646–650.

Akagi T, Pilkington SM, Varkonyi-Gasic E, Henry IM, Sugano SS, Sonoda M, Firl A, McNeilage MA, Douglas MJ, Wang T. 2019. Two Y chromosome-encoded genes determine sex in kiwifruit. bioRxiv:615666.

Anderson GJ. 1979. Dioecious Solanum species of hermaphroditic origin is an example of a broad convergence. Nature 282:836.

Anderson GJ, Anderson MK, Patel N. 2015. The ecology, evolution, and biogeography of dioecy in the genus Solanum: With paradigms from the strong dioecy in Solanum polygamum, to the unsuspected and cryptic dioecy in Solanum conocarpum. American Journal of Botany 102:471–486.

Anderson GJ, Levine DA. 1982. Three taxa constitute the sexes of a single dioecious species of Solanum. Taxon:667-672.

Anderson GJ, Symon DE. 1985. Extrafloral nectaries in Solanum. Biotropica:40-45.

Anderson GJ, Symon DE. 1989. Functional dioecy and andromonoecy in Solanum. Evolution 43:204–219.

Benjamini Y, Hochberg Y. 1995. Controlling the false discovery rate: a practical and powerful approach to multiple testing. Journal of the Royal Statistical Society. Series B (Methodological):289–300.

Bernardello LM, Anderson GJ. 1990. Karyotypic studies in Solanum section basarthrum (Solanaceae). American Journal of Botany 77:420–431.

Bosch M, Hepler PK. 2005. Pectin methylesterases and pectin dynamics in pollen tubes. The Plant Cell 17:3219–3226.

Bosch M, Hepler PK. 2006. Silencing of the tobacco pollen pectin methylesterase NtPPME1 results in retarded in vivo pollen tube growth. Planta 223:736–745.

Bräutigam K, Soolanayakanahally R, Champigny M, Mansfield S, Douglas C, Campbell MM, Cronk Q. 2017. Sexual epigenetics: gender-specific methylation of a gene in the sex determining region of Populus balsamifera. Scientific reports 7:45388.

Campbell MS, Law M, Holt C, Stein JC, Moghe GD, Hufnagel DE, Lei J, Achawanantakun R, Jiao D, Lawrence CJ. 2014. MAKER-P: a tool kit for the rapid creation, management, and quality control of plant genome annotations. Plant physiology 164:513–524.

Caporali E, Carboni A, Galli M, Rossi G, Spada A, Longo GM. 1994. Development of male and female flower in Asparagus officinalis. Search for point of transition from hermaphroditic to unisexual developmental pathway. Sexual Plant Reproduction 7:239–249.

Charlesworth B, Charlesworth D. 1978. A model for the evolution of dioecy and gynodioecy. The American Naturalist 112:975–997.

Charlesworth D. 2015. Plant contributions to our understanding of sex chromosome evolution. New Phytologist 208:52–65.

Charlesworth D. 2012. Plant sex chromosome evolution. Journal of experimental botany 64:405–420.

Charlesworth D. 2002. Plant sex determination and sex chromosomes. Heredity 88:94.

Cheung AY, Wu H-m. 2008. Structural and signaling networks for the polar cell growth machinery in pollen tubes. Annu. Rev. Plant Biol. 59:547–572.

Echeverría-Londoño S, Särkinen T, Fenton IS, Purvis A, Knapp S. 2020. Dynamism and context-dependency in diversification of the megadiverse plant genus Solanum (Solanaceae). Journal of Systematics and Evolution.

Ellegren H, Parsch J. 2007. The evolution of sex-biased genes and sex-biased gene expression. Nature Reviews Genetics 8:689.

Ellinghaus D, Kurtz S, Willhoeft U. 2008. LTRharvest, an efficient and flexible software for de novo detection of LTR retrotransposons. BMC bioinformatics 9:18.

Emms DM, Kelly S. 2015. OrthoFinder: solving fundamental biases in whole genome comparisons dramatically improves orthogroup inference accuracy. Genome biology 16:157.

Ezura K, Ji-Seong K, Mori K, Suzuki Y, Kuhara S, Ariizumi T, Ezura H. 2017. Genome-wide identification of pistil-specific genes expressed during fruit set initiation in tomato (Solanum lycopersicum). PloS One 12:e0180003.

Gaffe J, Tieman DM, Handa AK. 1994. Pectin methylesterase isoforms in tomato (Lycopersicon esculentum) tissues (effects of expression of a pectin methylesterase antisense gene). Plant Physiology 105:199–203.

Gou J-Y, Miller LM, Hou G, Yu X-H, Chen X-Y, Liu C-J. 2012. Acetylesterase-mediated deacetylation of pectin impairs cell elongation, pollen germination, and plant reproduction. The Plant Cell 24:50–65.

Grabherr MG, Russell P, Meyer M, Mauceli E, Alföldi J, Di Palma F, Lindblad-Toh K. 2010. Genome-wide synteny through highly sensitive sequence alignment: Satsuma. Bioinformatics 26:1145–1151.

Han MV, Thomas GW, Lugo-Martinez J, Hahn MW. 2013. Estimating gene gain and loss rates in the presence of error in genome assembly and annotation using CAFE 3. Molecular Biology and Evolution 30:1987–1997.

Han Y, Wessler SR. 2010. MITE-Hunter: a program for discovering miniature inverted-repeat transposable elements from genomic sequences. Nucleic acids research:gkq862.

Harkess A, Mercati F, Shan HY, Sunseri F, Falavigna A, Leebens-Mack J. 2015. Sex-biased gene expression in dioecious garden asparagus (asparagus officinalis). New Phytologist 207:883–892.

Harkess A, Zhou J, Xu C, Bowers JE, Van der Hulst R, Ayyampalayam S, Mercati F, Riccardi P, McKain MR, Kakrana A. 2017. The asparagus genome sheds light on the origin and evolution of a young Y chromosome. Nature communications 8:1279.

Henry IM, Akagi T, Tao R, Comai L. 2018. One hundred ways to invent the sexes: Theoretical and observed paths to dioecy in plants. Annual review of plant biology 69:553–575.

Jiang J, Zhang Z, Cao J. 2013. Pollen wall development: the associated enzymes and metabolic pathways. Plant biology 15:249–263.

Kazama Y, Ishii K, Aonuma W, Ikeda T, Kawamoto H, Koizumi A, Filatov DA, Chibalina M, Bergero R, Charlesworth D. 2016. A new physical mapping approach refines the sex-determining gene positions on the Silene latifolia Y-chromosome. Scientific reports 6:18917.

Kim D, Langmead B, Salzberg SL. 2015. HISAT: a fast spliced aligner with low memory requirements. Nature methods 12:357.

Knapp S, Bohs L, Nee M, Spooner DM. 2004. Solanaceae—a model for linking genomics with biodiversity. Comparative and functional genomics 5:285–291.

Korneliussen TS, Albrechtsen A, Nielsen R. 2014. ANGSD: analysis of next generation sequencing data. BMC bioinformatics 15:356.

Levine DA, Anderson GJ. 1986. Evolution of Dioecy in an American Solanum. In: D’Arcy WG, editor. Solanaceae: Biology and Systematics. New York: Columbia University Press. p. 264–273.

Li H, Handsaker B, Wysoker A, Fennell T, Ruan J, Homer N, Marth G, Abecasis G, Durbin R. 2009. The sequence alignment/map format and SAMtools. Bioinformatics 25:2078-2079.

Li S-F, Zhang G-J, Yuan J-H, Deng C-L, Gao W-J. 2016. Repetitive sequences and epigenetic modification: inseparable partners play important roles in the evolution of plant sex chromosomes. Planta 243:1083–1095.

Liao Y, Smyth GK, Shi W. 2013. featureCounts: an efficient general purpose program for assigning sequence reads to genomic features. Bioinformatics 30:923–930.

Marçais G, Kingsford C. 2011. A fast, lock-free approach for efficient parallel counting of occurrences of k-mers. Bioinformatics 27:764–770.

Martin A, Troadec C, Boualem A, Rajab M, Fernandez R, Morin H, Pitrat M, Dogimont C, Bendahmane A. 2009. A transposon-induced epigenetic change leads to sex determination in melon. Nature 461:1135.

McDonnell AJ, Wetreich HB, Cantley JT, Jobson P, Martine CT. 2019. Solanum plastisexum, an enigmatic new bush tomato from the Australian Monsoon Tropics exhibiting breeding system fluidity. PhytoKeys 124:39.

Micheli F. 2001. Pectin methylesterases: cell wall enzymes with important roles in plant physiology. Trends in plant science 6:414–419.

Mollet J-C, Leroux C, Dardelle F, Lehner A. 2013. Cell wall composition, biosynthesis and remodeling during pollen tube growth. Plants 2:107–147.

Murase K, Shigenobu S, Fujii S, Ueda K, Murata T, Sakamoto A, Wada Y, Yamaguchi K, Osakabe Y, Osakabe K. 2017. MYB transcription factor gene involved in sex determination in Asparagus officinalis. Genes to Cells 22:115–123.

Nag A, Jack T. 2010. Sculpting the flower; the role of microRNAs in flower development. In. Current opics in Developmental Biology: Elsevier. p. 349–378.

Oh SA, Park KS, Twell D, Park SK. 2010. The SIDECAR POLLEN gene encodes a microspore-specific LOB/AS2 domain protein required for the correct timing and orientation of asymmetric cell division. The Plant Journal 64:839–850.

Otto SP, Pannell JR, Peichel CL, Ashman T-L, Charlesworth D, Chippindale AK, Delph LF, Guerrero RF, Scarpino SV, McAllister BF. 2011. About PAR: the distinct evolutionary dynamics of the pseudoautosomal region. Trends in Genetics 27:358–367.

Ou S, Jiang N. 2018. LTR_retriever: A Highly Accurate And Sensitive Program For Identification Of LTR Retrotransposons. Plant Physiology 176:1410–1422.

Quinlan AR, Hall IM. 2010. BEDTools: a flexible suite of utilities for comparing genomic features. Bioinformatics 26:841–842.

Renner SS. 2014. The relative and absolute frequencies of angiosperm sexual systems: dioecy, monoecy, gynodioecy, and an updated online database. American Journal of Botany 101:1588–1596.

Rice WR. 1987. The accumulation of sexually antagonistic genes as a selective agent promoting the evolution of reduced recombination between primitive sex chromosomes. Evolution 41:911–914.

Robinson MD, McCarthy DJ, Smyth GK. 2010. edgeR: a Bioconductor package for differential expression analysis of digital gene expression data. Bioinformatics 26:139–140.

Robinson MD, Oshlack A. 2010. A scaling normalization method for differential expression analysis of RNA-seq data. Genome biology 11:R25.

Särkinen T, Bohs L, Olmstead RG, Knapp S. 2013. A phylogenetic framework for evolutionary study of the nightshades (Solanaceae): a dated 1000-tip tree. BMC Evolutionary Biology 13:214.

Schmieder R, Edwards R. 2011. Fast identification and removal of sequence contamination from genomic and metagenomic datasets. PloS One 6:e17288.

Shi J, Cui M, Yang L, Kim Y-J, Zhang D. 2015. Genetic and biochemical mechanisms of pollen wall development. Trends in plant science 20:741–753.

Simão FA, Waterhouse RM, Ioannidis P, Kriventseva EV, Zdobnov EM. 2015. BUSCO: assessing genome assembly and annotation completeness with single-copy orthologs. Bioinformatics:btv351.

Smit A, Hubley R, Green P. 2015. RepeatMasker Open-4.0. 2013–2015. Institute for Systems Biology.

Song Y, Ma K, Ci D, Zhang Z, Zhang D. 2013. Sexual dimorphism floral microRNA profiling and target gene expression in andromonoecious poplar (Populus tomentosa). PLoS One 8:e62681.

Symon D editor. Linnean Society symposium series. 1979.

Symon DE. 1981. A revision of the genus Solanum in Australia. Journal of the Adelaide Botanic Gardens 4:1–367.

The Tomato Genome Consortium. 2012. The tomato genome sequence provides insights into fleshy fruit evolution. Nature 485:635–641.

Thomas GW, Hahn MW. 2019. Referee: reference assembly quality scores. Genome biology and evolution 11:1483–1486.

Torres MF, Mathew LS, Ahmed I, Al-Azwani IK, Krueger R, Rivera-Nuñez D, Mohamoud YA, Clark AG, Suhre K, Malek JA. 2018. Genus-wide sequencing supports a two-locus model for sex-determination in Phoenix. Nature communications 9:3969.

Vurture GW, Sedlazeck FJ, Nattestad M, Underwood CJ, Fang H, Gurtowski J, Schatz MC. 2017. GenomeScope: fast reference-free genome profiling from short reads. Bioinformatics 33:2202–2204.

Wang J, Na J-K, Yu Q, Gschwend AR, Han J, Zeng F, Aryal R, VanBuren R, Murray JE, Zhang W. 2012. Sequencing papaya X and Yh chromosomes reveals molecular basis of incipient sex chromosome evolution. Proceedings of the National Academy of Sciences of the United States of America 109:13710–13715.

Whalen MD, Costich DE. 1986. Andromonoecy in solanum. Solanaceae: biology and systematics:284–302.

Wu H-C, Bulgakov VP, Jinn T-L. 2018. Pectin methylesterases: cell wall remodeling proteins are required for plant response to heat stress. Frontiers in Plant Science 9.

Wu M, Kostyun J, Moyle L. 2019. Genome sequence of Jaltomata addresses rapid reproductive trait evolution and enhances comparative genomics in the hyper-diverse Solanaceae. Genome Biology and Evolution 11:335–349.

Zavada MS, Anderson GJ. 1997. The wall and aperture development of pollen from dioecious Solanum appendiculatum: What is inaperturate pollen? Grana 36:129–134.

Zhang GY, Feng J, Wu J, Wang XW. 2010. BoPMEI1, a pollen-specific pectin methylesterase inhibitor, has an essential role in pollen tube growth. Planta 231:1323–1334.

Zhu J, Zhang G, Chang Y, Li X, Yang J, Huang X, Yu Q, Chen H, Wu T, Yang Z. 2010. AtMYB103 is a crucial regulator of several pathways affecting Arabidopsis anther development. Science China Life Sciences 53:1112–1122.

Zimin AV, Puiu D, Luo M-C, Zhu T, Koren S, Yorke JA, Dvorak J, Salzberg S. 2017. Hybrid assembly of the large and highly repetitive genome of Aegilops tauschii, a progenitor of bread wheat, with the mega-reads algorithm. Genome Research 29:2669–2677.

